# Structural basis of auxin binding and transport by *Arabidopsis thaliana* AUX1

**DOI:** 10.1101/2025.05.27.656222

**Authors:** Dan Jing, Fang Kong, Xiaoli Lu, Gaoxingyu Huang, Jing Huang, Haolin Wang, Yigong Shi, Chengcheng Wang

## Abstract

Indole-3-acetic acid (IAA), the major form of auxin, is essential for plant growth. Auxin-resistant 1 (AUX1), the first identified auxin importer, plays a crucial role in polar auxin transport (PAT). Here we present cryo-EM structures of *Arabidopsis thaliana* AUX1 in the IAA-free and IAA-bound states. AUX1 exists as a monomer that contains 11 transmembrane helixes (TMs). TMs 1-5 and 6-10 constitute the two halves of a classic LeuT-fold, and TM11 interacts with both halves at the interface. In the IAA-bound state, IAA is specifically recognized in a central pocket formed by TM1, TM3, TM6, and TM8. In the presence of IAA, TM1 and TM6 undergo marked conformational changes that are critical for IAA transport. His249 stands out to be a key residue for substrate uptake and release. Our structures reveal the molecular basis for AUX1-mediated IAA binding and transport.

**Significance:** Auxin-resistant 1 (AUX1) is a major auxin influx transporter, playing critical regulatory roles in callus growth, vascular patterning, leaf phyllotactic patterning, root aerial development, and apical hook formation. However, its working mechanism remains unclear. In this study, we report two structures of IAA-bound and IAA-free AUX1, elucidating its substrate recognition mechanism and proposing a potential substrate release model for AUX1. These findings offer important insights into the molecular mechanisms of AUX1-mediated IAA binding and transport, and lay the foundation for future structure-based functional studies of the AUX1/LAX family and the application of auxin analogs in agriculture.

## Introduction

Auxin participates in almost all aspects of plant growth and development, including cell augmentation, division, differentiation, signal transmission, and environmental adaptability (1). Indole-3-acetic acid (IAA), the most predominant naturally occurring auxin, is widely distributed in green plants (2, 3). IAA is primarily synthesized in the shoot apical meristem, leaf primordia, and other growing tissues, and is transported to the target tissues either through the polar auxin transport (PAT) stream or via the bulk flow in the phloem (4, 5). At the cellular level, PAT involves both passive membrane diffusion of the protonated form (IAAH) and active transport of the ionized form (IAA^-^) mediated by specific carrier proteins.

Normally, with the extracellular apoplast pH ranging from 5.0 to 5.5, only 15-25% of auxin (pKa ∼ 4.75) is protonated and capable of passive diffusion, whereas the majority requires carriers-mediated cellular entry (1). Several auxin transporters have been identified, including the auxin-resistant 1 (AUX1) and like-AUX1 (LAX) family, the PIN-FORMED (PIN) family, and certain members of the P-glycoprotein/ATP-binding cassette B4 (PGP/ABCB) family (6). Among these, AUX1 is a proton-auxin symporter that belongs to the amino acid/auxin permease (AAAP) family (7, 8).

In *Arabidopsis thaliana* (*A. thaliana*), AUX1 is localized to the plasma membrane and serves as a key player in auxin uptake. The AUX1 protein is mainly expressed in the phloem, vascular cylinder, lateral root cap, and epidermal cells of the root tip (9, 10). The basipetal auxin transport defects associated with AUX1 mutations can lead to decreased seed germination, reduced root growth, disrupted root gravitropism, or impaired lateral root development (7, 9, 11-14). AUX1 is also implicated in the regulation of various developmental processes, including callus growth, vascular patterning, leaf phyllotactic patterning, root air development, and apical hook formation (15-19).

Despite its well-established importance in plant growth and development, the AUX1/LAX family is yet to be structurally characterized. Recently, the cryo-electron microscopy (cryo-EM) structures of the PIN family members were reported, offering important mechanistic insights into auxin efflux (20-22). In contrast, the molecular basis of auxin influx remains unclear. Here, we present the IAA-free and IAA-bound structures of the *A. thaliana* AUX1, which provide mechanistic insights into substrate recognition and transport.

## Results

### Overall structure of AUX1

In *A. thaliana*, the AUX1/LAX family comprises four members: AUX1, LAX1, LAX2, and LAX3, which share 75–80% similarity at the amino acid sequence level (*SI Appendix*, Fig. S1). Their auxin influx activities have been demonstrated in heterologous expression systems using human HEK293 cells, U2OS cells, yeasts, or *Xenopus laevis* oocytes (23-26). We sought to determine the structure of AUX1, the founding member of this family that has been subjected to extensive investigations. First, we established a radioactive [^3^H]IAA transport system to examine the activity of the *A. thaliana* AUX1 (Fig. 1*A*). The HEK293T cells with AUX1 expression showed a higher level of [^3^H]IAA accumulation compared to the control cells (empty vector without AUX1) (Fig. 1*B*). Furthermore, each of the three auxin influx inhibitors 1-naphthoxyacetic acid (1-NOA), 3-chloro-4-hydroxyphenylacetic acid (CHPAA), and 2-naphthoxyacetic acid (2-NOA) reduced the uptake of [^3^H]IAA in AUX1-expressing HEK293T cells (Fig. 1*B*). These results confirm AUX1 as a direct mediator of active auxin influx.

**Figure. 1.**
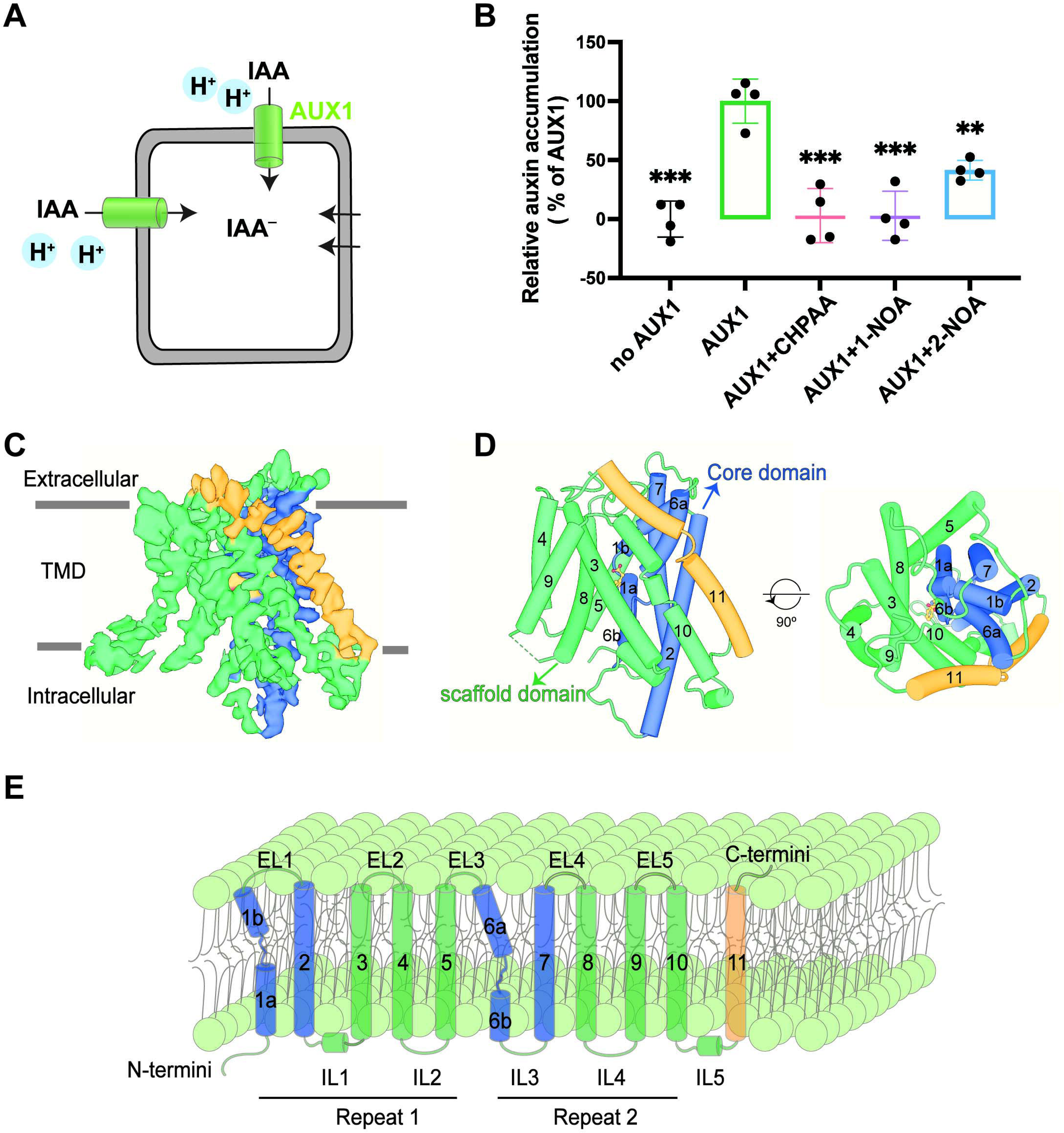
Overall structure of AUX1. (A) A schematic illustration of the IAA uptake assay. IAA^-^ represents the deprotonated form. IAA^-^ and protons (H^+^) are thought to be taken up by AUX1 from the extracellular side and transported into the cell. In the actual assay, IAA is labeled by the isotope ^3^H. (B) AUX1 mediates IAA import in HEK293T cells. In addition, uptake is reduced upon incubation with the auxin influx-specific inhibitors CHPAA, 1-NOA, and 2-NOA. Shown here are results of relative auxin accumulation. The result for cells with AUX1 expression is normalized as 100%. Data are mean ± s.d.; n=4 (***P < 0.001, **P < 0.01, Two-tailed unpaired t-test). Independent experiments have been repeated for four times with similar results. (C) Cryo-EM reconstruction of AUX1 treated with IAA. The EM map, at an overall resolution of 3.5 Å, is contoured at level 1.1 and colored by domain. (D) Overall structure of the AUX1-IAA complex. Two perpendicular views are shown. TMs 1, 2, 6, and 7 form the core domain (colored blue). TMs 3-5 and TMs 8-10 constitute the scaffold domain (colored green). TM11 (colored yellow) binds the interface between the core and scaffold domains. (E) The topology diagram of AUX1. 11 TMs and the intervening loops are labeled. EL: extracellular loop; IL: intracellular loop. All structural images in this paper were generated in UCSF ChimeraX (46, 47).

The *A. thaliana* AUX1 protein has a modest molecular weight of ∼54 kD and lacks sufficiently large soluble domains, both posing serious challenges for its cryo-EM analysis. To facilitate the structure determination of AUX1, we screened multiple recombinant expression systems to increase protein production and tested various detergents to optimize preparation of the cryo-samples. After numerous trials, we were able to express AUX1 in HEK293F cells and extracted the recombinant protein using 0.002% (w/v) lauryl maltose neopentyl glycol (LMNG) plus 0.0002% (w/v) cholesteryl hemisuccinate tris salt (CHS) (*SI Appendix*, Fig. S2 *A* and *B*). An excess amount of the substrate IAA was added during cryo-sample preparation. We collected two sets of cryo-EM data on a Titan Krios electron microscope (*SI Appendix*, Fig. S2*C*).

We started with a set of 3,837 micrographs to generate a good initial EM reconstruction, which was used for further data processing (*SI Appendix*, Fig. S2*D*). Finally, relying on 12,663 high-quality micrographs, we obtained a 3D reconstruction of the AUX1-IAA complex at an average resolution of 3.5 Å (Fig. 1*C* and *SI Appendix*, Fig. S2 *D*-*G*). The EM map was of sufficient quality to support atomic modeling of 415 residues (Fig. S3).

The structure of AUX1 contains 11 transmembrane helices (TMs 1-11), with the N- and C-termini located on the cytosolic and extracellular sides of the cell membrane, respectively (Fig. 1*C*-*E*). TMs 1-10 of AUX1 adopt a typical LeuT-fold, wherein TMs 1-5 and TMs 6-10 form two inverted structural repeats (*SI Appendix*, Fig.S4). TMs 3-5 and TMs 8-10 constitute the “scaffold domain”; TMs 1, 2, 6, and 7 stack together to form the “core domain” (Fig. 1*D*). The structure immediately suggests a substrate transport path between the scaffold domain and the core domain (Fig. 1*D*). AUX1 has an additional, extended transmembrane helix TM11. With a kink in the center, TM11 wraps one side of the transporter with a tilt angle of ∼ 45° to the membrane norm (Fig. 1 *C* and *D*). TM11 binds the interface between the two structural repeats, likely stabilizing the overall structure of AUX1.

### IAA recognition by AUX1

As in all LeuT-fold transporters, TM1 and TM6 of AUX1 are discontinuous and each comprises two half helices: TM1a/TM1b and TM6a/TM6b (Fig. 1 *D* and *E*). A relatively narrow, solvent-accessible vestibule extends from the intracellular side to the predicted substrate-binding pocket, suggesting a substrate-bound, inward-facing conformation for AUX1 (Fig. 2*A*). After completion of atomic modeling for AUX1, an extra EM density lobe remains unaccounted for in the substrate-binding pocket (Fig. 2*B*). We tentatively assigned an IAA molecule to this density lobe (Fig. 2*C* and *SI Appendix*, Fig. S3*B*). This assignment is consistent with the AlphaFold3-predicted IAA binding mode for AUX1 (27) (*SI Appendix*, Fig. S5*A*). However, the AlphaFold3-predicted main chain around His249 is inconsistent with our EM density map (*SI Appendix*, Fig. S5*B*); in our final model, this part of the main chain closely matches the EM density, placing the imidazole side chain of His249 in contact with the indole ring of IAA (*SI Appendix*, Fig. S5*C*).

**Figure. 2.**
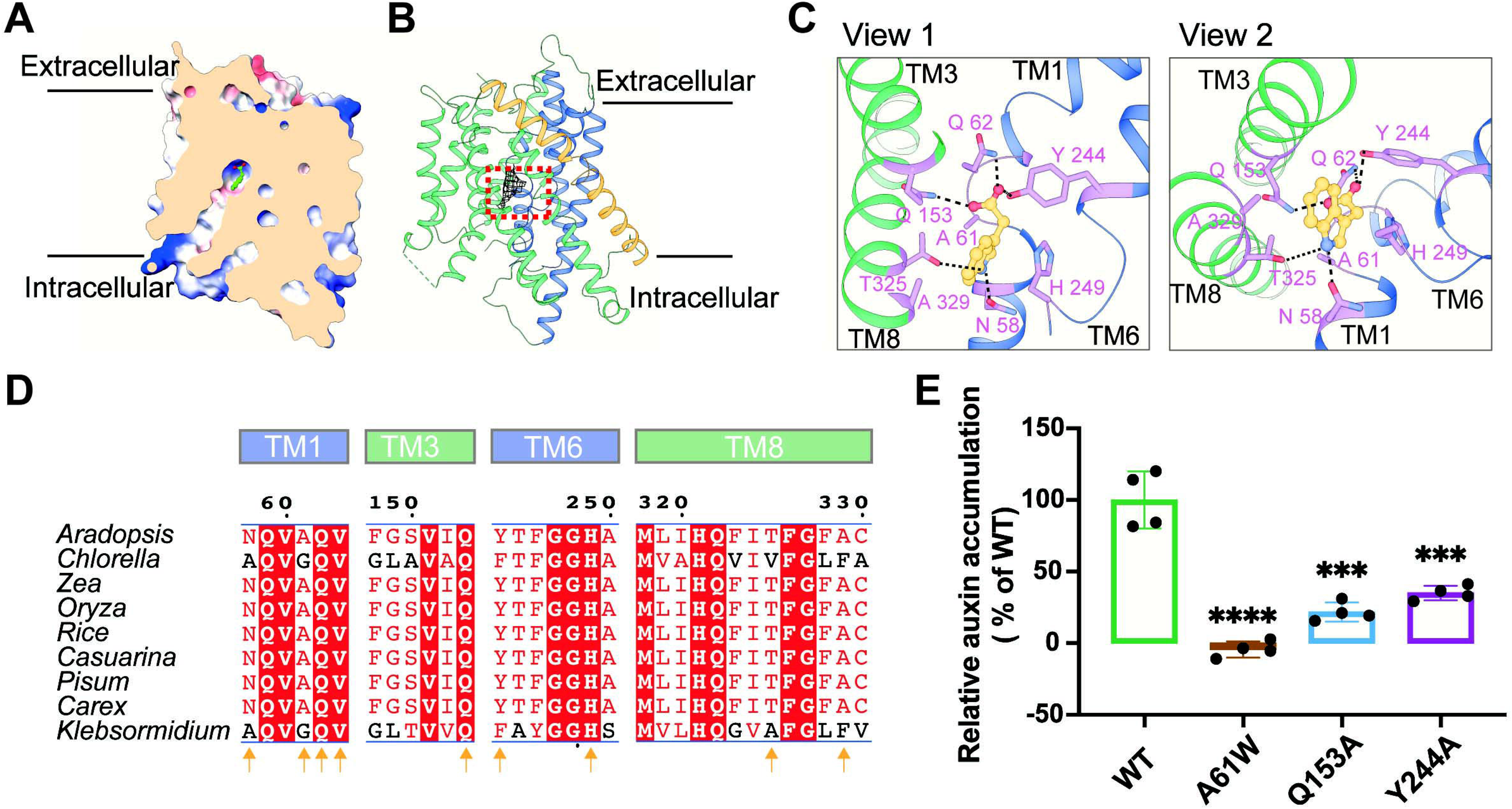
IAA is recognized in an inward-facing conformation of AUX1. (A) The structure of AUX1 bound to IAA exists in an inward-facing conformation. Negative and positive charges are colored red and blue, respectively. Shown here is a cut-open view of the surface electrostatic potential of AUX1 that is calculated in UCSF ChimeraX. (B) IAA is bound at the center of AUX1. The EM density of the substrate IAA, contoured at level 0.86, is shown as grey mesh. AlphFold3 (27) was employed to facilitate assignment of IAA. (C) Coordination of IAA by residues from TM1, TM3, TM6, and TM8. Two perpendicular views are shown. IAA is shown as yellow ball-and-stick, and IAA-coordinating residues are highlighted as pink sticks. Putative H-bonds are represented by black, dashed lines. (D) IAA-binding residues are highly conserved across species. Shown here is an alignment of the sequence elements that constitute the IAA-binding pocket. Residues that may recognize IAA are marked by yellow arrows. (E) Mutation of IAA-binding residues in AUX1 results in compromised auxin accumulation in HEK293T cells. Shown here are results of [^3^H]IAA uptake assays. The result for cells with WT AUX1 expression is normalized as 100%. All experiments were performed in triplicates. Data are mean ± s.d.; n=4 (****P < 0.0001, ***P < 0.001, Two-tailed unpaired t-test). Independent experiments have been repeated for three times with similar results.

The IAA-binding pocket is formed by TM1, TM3, TM6, and TM8 of AUX1. IAA is likely recognized through both hydrogen bonds (H-bonds) and van der Waals contacts (Fig. 2*C*). The carboxyl group of IAA appears to be H-bonded with the side chains of Tyr244, Gln62, and Gln153. The amide group of the imidazole ring of IAA may form H-bonds with Asn58 and Thr325 (Fig. 2*C*), and the indole group likely make van der Waals contacts to Ala61, His249, and Ala329. These residues are highly conserved among plants (Fig. 2*D*).

To corroborate the structural assignment, we generated several AUX1 mutants, each containing a missense mutation that targets a conserved coordinating residue, and measured their IAA transport activities using the aforementioned [^3^H]IAA influx assay in HEK293T cells. These mutant AUX1 proteins were expressed at a similar level as the wild-type (WT) AUX1 and localized at the plasma membrane (*SI Appendix*, Fig. S6). Consistent with our structural observations, Ala substitutions of Gln153 and Tyr244, expected to disrupt the H-bonds, result in dramatically reduced transport activities of AUX1 (Fig. 2*E*). The A61W mutation also substantially attenuated the transport activity, suggesting that a large side chain might hinder IAA binding. Four AUX1 mutants (T245A, G247E, A250T, and F145A) displayed severely compromised protein expression levels, preventing assessment of their effects on transport activity.

### Conformational changes in response to IAA binding

To gain insights into substrate transport, we set out to determine the cryo-EM structure of IAA-free AUX1. In our hands, IAA-free AUX1 repeatedly defied cryo-sample preparation. To improve the sample quality, we inserted a BRIL domain into the loop (named IL1 for the intracellular loop 1) connecting TM2 and TM3 (*SI Appendix*, Fig. S7*A*). The resulting AUX1_IL1_BRIL fusion protein exhibited the same transport activity as the WT protein (*SI Appendix*, Fig. S7*B*). We then incubated purified AUX1_IL1_BRIL with an anti-BRIL Fab fragment (Fab) and an anti-Fab nanobody (Nb), purified the complex (*SI Appendix*, Fig. S8*A*), and subjected the complex to EM analysis (*SI Appendix*, Fig. S8*B*).

Relying on 20,418 micrographs, we obtained a final reconstruction of the IAA-free AUX1_IL1_BRIL/Fab/Nb complex at an average resolution of 3.97 Å (*SI Appendix*, Fig. S8*C*-*F*, and S9). In the EM map, the BRIL domain and its associated Fab and nanobody are clearly identified (*SI Appendix*, Fig. S9 *A* and *C*); but these structural elements are separated from the IAA-free AUX1 and will not be discussed further in this manuscript.

We built an atomic model for the IAA-free AUX1 (*SI Appendix*, Fig. S9*D*). The EM map for TM6b was relatively weak; this part of the model (residues 247-254) was generated mainly on the basis of AlphaFold2 prediction (AF-Q96247-F1-v4) (28). We superimposed the final atomic coordinates of the IAA-free AUX1 with those of the IAA-bound AUX1 (Fig. 3*A*). Compared to the IAA-bound state, TM1 and TM6, which play an important role in IAA binding, display marked conformational shifts in the IAA-free state. Importantly, no EM densities are observed near the IAA binding site in the IAA-free AUX1, supporting the assignment of IAA in the previous EM reconstruction of the IAA-bound AUX1.

**Figure. 3.**
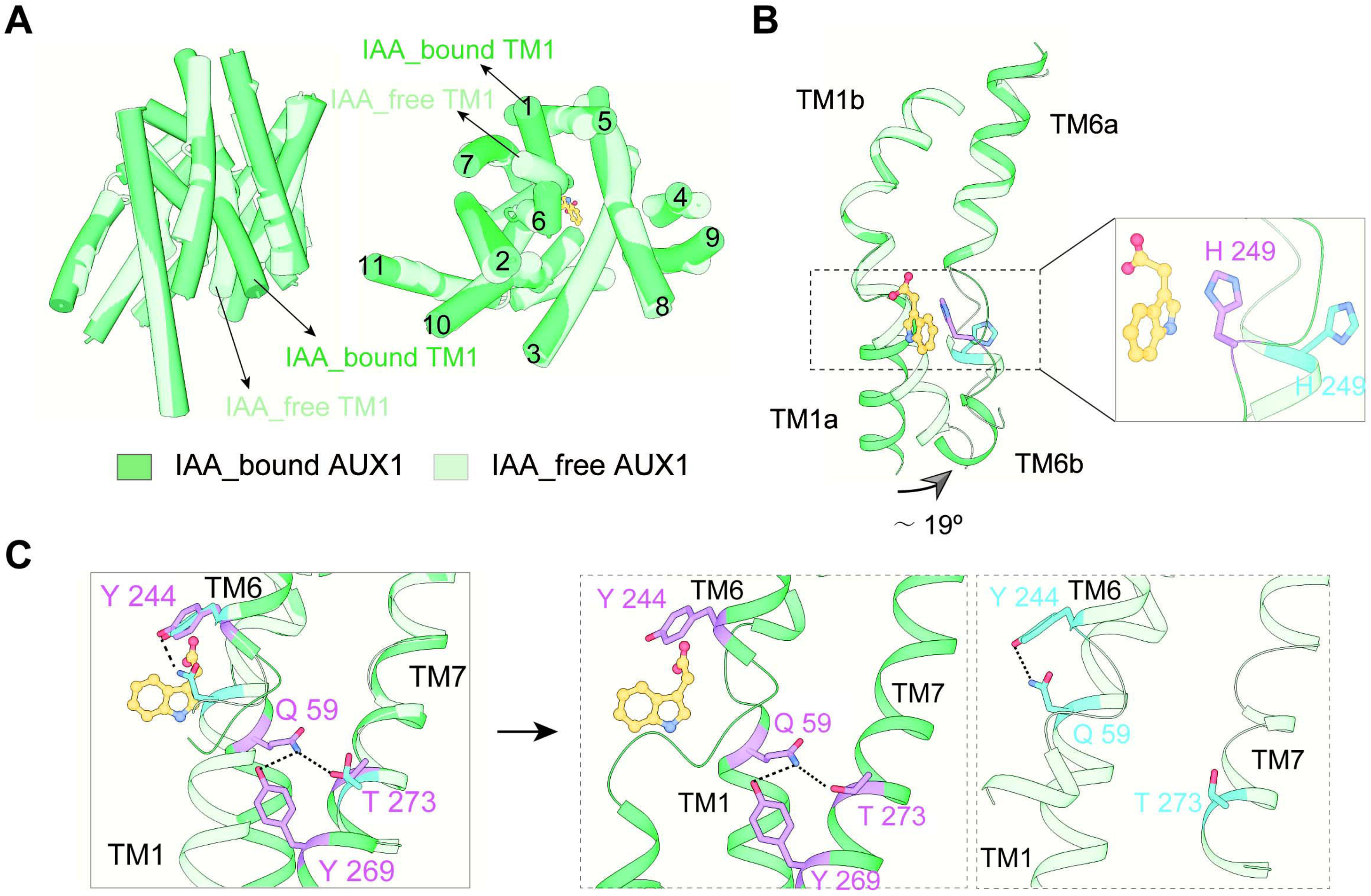
Conformational changes between IAA-bound and IAA-free AUX1. (A) Comparison of the overall structures of IAA-bound (green) and IAA-free (light green) AUX1. These two structures can be superimposed with a root mean squared deviation (RMSD) of 0.933 Å over 316 Cα atoms. A notable change occurs to TM1. (B) A close-up view on the structural comparison of TM1 and TM6 between the IAA-bound and IAA-free states. TM1a is rotated by about 19 degrees to accommodate the substrate. The inset shows the configurational switch of the side chain of His249 in response to IAA binding. (C) A potential role of Gln59 in IAA binding and release. The left panel shows a comparison of the local structural elements surrounding Gln59 between the IAA-bound and IAA-free states. The middle and right panels show these elements in the IAA-bound and IAA-free states, respectively. In the IAA-free state, Gln59 interacts with Tyr244 on TM6a (right panel). In the IAA-bound state, Gln59 shifts its side chain to make potential H-bonds with Thr273 and Try 269 on TM7 (middle panel).

A closer examination of the structure overlay reveals a rotation of approximately 19° for TM1a and a configurational switch of His249 on TM6 to accommodate the substrate (Fig. 3*B*). Gln59 on TM1a in the IAA-free state appears to interact with Tyr244 on TM6a. Importantly, this configuration of Gln59 would clash with the IAA molecule. In the IAA-bound state, Gln59 interacts with Thr273 and Try269 on TM7 through H-bonds (Fig. 3*C*).

## Discussion

### IAA binding and release

From embryogenesis to fruit formation, various stages of plant development are controlled by the differential distribution of auxin across cells and tissues (29). Here we report the overall structure and substrate recognition of AUX1, the predominant auxin importer in *A. thaliana*. Comparison of the inward-facing IAA-bound and IAA-free structures suggests a critical role of the broken helices TM1 and TM6 in substrate binding and release (Fig. 4). It is noted that similar substrate binding sites have been identified in LeuT-fold AAAP family transporters (30, 31), but the coordinating residues and TMs vary, consistent with their respective substrate specificity (*SI Appendix*, Fig. S10). For instance, MhsT is a bacterial transporter for hydrophobic amino acids; recognition of tryptophan or 4-fluoro-L-phenylalanine has been documented (PDB codes: 4US3 and 6YU4) (31, 32). Although tryptophan serves as the biosynthetic precursor of IAA, the details for their respective recognition by MhsT and AUX1 are different (*SI Appendix*, Fig. S10), consistent with prior evidence that tryptophan fails to inhibit AUX1-mediated transport activity (33).

**Figure. 4.**
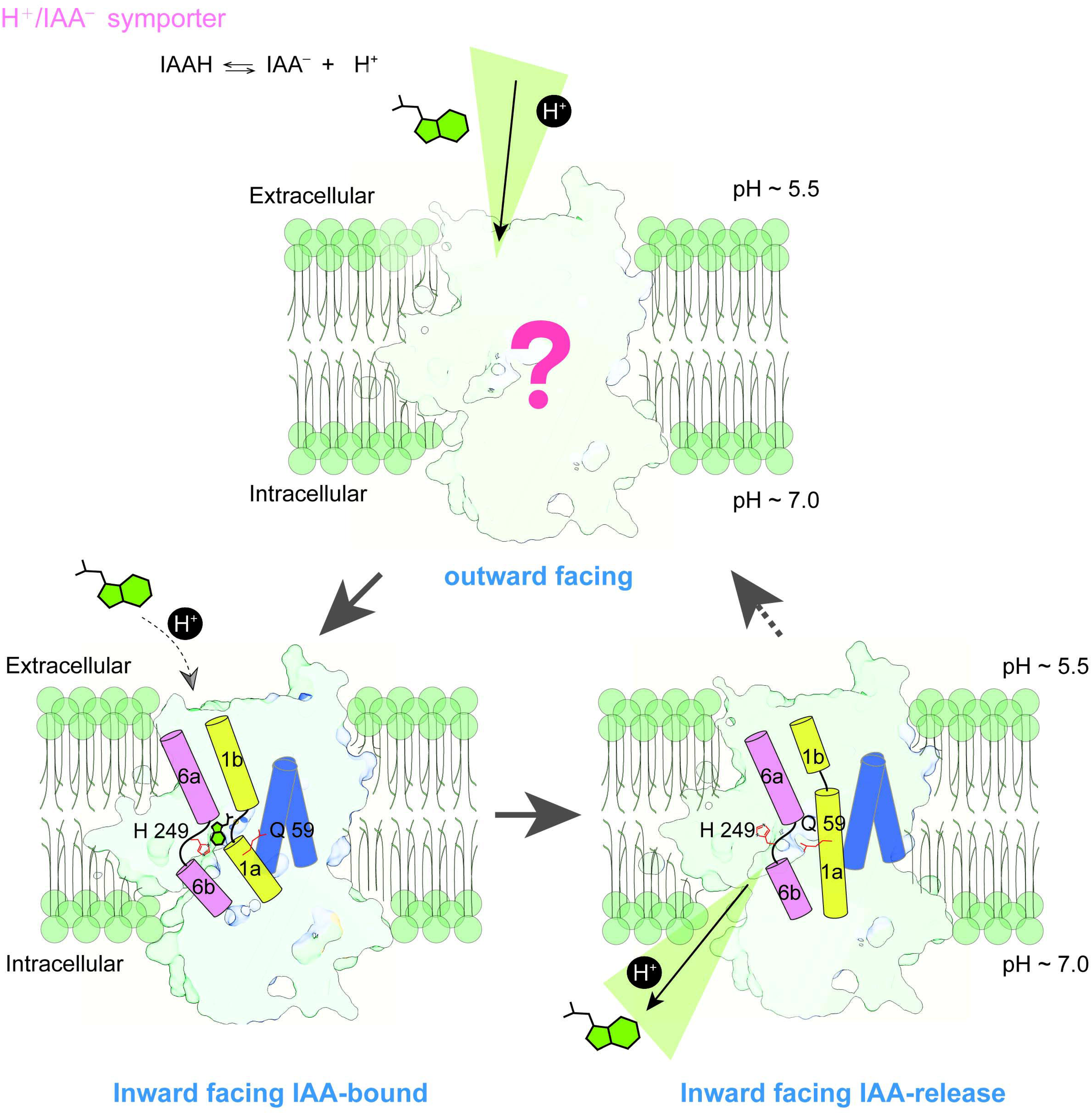
A working model of substrate binding and release for AUX1. With a pKa of about 4.75, the vast majority of IAA remains in the deprotonated state (IAA^-^) in the extracellular environment. In its outward-facing state, AUX1 specifically binds IAA^-^. This process is likely facilitated by protonation of His249, thereby satisfying the H^+^ symport mechanism. Next, AUX1 changes to the inward-facing conformation, exposing the IAA-binding pocket to the intracellular pH of about 7.4. Under this condition, His249 likely becomes deprotonated, causing local conformational changes. Additionally, the binding and release of IAA induce a coordinated conformational shift in TM1a, which pushes Gln59 into the binding pocket to cause steric hindrance, thereby potentiating substrate dissociation.

In AUX1, His249 appears to play a key role, directly contributing to IAA recognition in the IAA-bound state, but moves away from the IAA-binding pocket in the IAA-free state. Additionally, the binding and release of IAA induce a coordinated conformational shift in TM1a, which pushes Gln59 into the binding pocket to cause steric hindrance, thereby potentiating substrate dissociation. The swinging of TM1a has also been observed in other LeuT-fold transporters, such as LeuT, SERT, LAT1, and MntH (34). Notably, His249 and Gln59 are invariable among all AUX1/LAX family members (*SI Appendix*, Fig. S1), suggesting a conserved mechanism for substrate binding and release.

We have yet to capture the outward-facing conformation of AUX1, which would be crucial for understanding the transport mechanism of IAA. Further structural studies are required to reveal the complete transport cycle of AUX1 (Fig. 4).

### Proton symport mechanism

The protonation/deprotonation of His249 (pKa ∼ 6.0), which is located in the IAA-binding pocket, may take part in the AUX1-mediated co-transporting proton and IAA uptake (35-37) (Fig. 4). With a pKa of about 4.75, the vast majority of IAA remains in the deprotonated state (IAA^-^) in the extracellular environment (pH ∼5.0-5.5). In its outward-facing state, AUX1 specifically binds IAA^-^ and His249 is predominantly protonated. Importantly, IAA^-^ binding is likely facilitated by protonation of His249, thereby satisfying the H^+^ symport mechanism. Next, AUX1 changes to the inward-facing conformation, exposing the IAA-binding pocket to the intracellular pH of about 7.4. Under such pH, His249 should be deprotonated when exposed to the cytosol, causing local conformational rearrangements.

To verify this hypothesis, we examined the IAA binding affinity of AUX1 and its mutant protein AUX1^H249A^ at different pH values using surface plasmon resonance (SPR). The IAA binding affinity of WT AUX1 at pH 5.5 (mimicking extracellular milieu) is an order of magnitude higher than that at pH 7.4 (mimicking the cytosolic environment) (*SI Appendix*, Figure S11 *A* and *B*), consistent with previous reports (35). Notably, compared to WT AUX1, the IAA binding affinity of AUX1^H249A^ remains largely unchanged at pH 7.4, but is drastically decreased by more than two orders of magnitude at pH 5.5 (*SI Appendix*, Fig. S11 *C* and *D*). The pH-dependent binding affinities for IAA may allow AUX1 to avidly bind IAA on the extracellular side and release it on the cytoplasmic side (*SI Appendix*, Fig. S11*E*). These results establish the significance of His249 protonation. We also attempted to determine the structure of AUX1 bound to IAA at pH 5.5 but failed to gain a high-resolution structure.

We further explored the potential role of His249 protonation by performing all-atom molecular dynamics (MD) simulations, with His249 in three protonation states (N^ε2^-protonated, N^δ1^-protonated, and doubly protonated) (*SI Appendix*, Fig. S12). In the absence of IAA^-^, His249 in the N^ε2^-protonated state (HSE) appears to be most stable (*SI Appendix*, Fig. S12*A*, left panel). In the presence of IAA^-^, His249 in the doubly protonated (HSP) state becomes more stable (*SI Appendix*, Fig. S12*A*, right panel). In accordance with this analysis, IAA is also most stable in the HSP state (*SI Appendix*, Fig. S12*B*). These findings support a model where protonation and deprotonation of the N^δ1^ atom of His249 are coupled to the binding and release of IAA, respectively.

### Structure-based mutational analysis

Based on sequence alignment (*SI Appendix*, Fig. S1 and S14), we mapped a subset of missense mutations that have been reported to cause plant growth defects onto the AUX1 structure (*SI Appendix*, Fig. S13) (10). The vast majority of these mutations are mapped to the packing interface among the TMs, likely damaging the structural stability of AUX1. Nonetheless, a few mutations affect residues in the IAA binding pocket and may negatively influence substrate recognition.

In conclusion, our study offers important insights into the molecular mechanisms of AUX1-mediated IAA binding and transport, and sets the foundation for future structure-based functional studies of AUX1/LAX family and application of auxin analogs in agriculture.

## Materials and Methods

### Protein expression and purification

The cDNA sequences of full-length *Arabidopsis thaliana* AUX1 are publicly available at Uniprot (https://www.uniprot.org) under the accession code of Q96247 and were cloned into the pCAG vector. For the transport assay, the full-length AUX1 was fused with an enhanced green fluorescent protein (EGFP), a FLAG tag (DYKDDDDK), and a 10 × His tag (HHHHHHHHHH) at its N-terminus. For IAA-bound AUX1 structure determination, a FLAG tag and a 10 × His tag were fused to the N-terminus of AUX1. For structure determination of IAA-free AUX1, a FLAG tag and 8 × His tag were fused to the N-terminus of AUX1, and an extra BRIL sequence (1) was introduced after codon 116 in AUX1 (AUX1_IL1_BRIL). Point mutations were generated using a standard two-step PCR strategy. The heavy and light chain sequences of anti-BRIL Fab (1) were separately cloned into pCAG vector that bears an IGK signal sequence. A 6 × His tag was fused to the C-terminus of the heavy chain. All constructs were verified by DNA sequencing.

For protein expression, HEK293F cells (Sino Biological) were cultured in SMM 293T-I medium (Sino Biological Inc.) at 37 ℃ with 5% CO_2_ in a Multitron-Pro shaker (Infors). To monitor the protein expression patterns of AUX1 in HEK293F/HEK293T cells, AUX1 plasmids fused with an N-terminal EGFP were transiently transfected into HEK293F/HEK293T cells using PEI (YEASEN). After culturing for 48 hours at 37 °C, the GFP fluorescence was observed using a Nikon Spinning Disk Field Scanning Confocal Systems (Nikon) with an excitation wavelength of 488 nm.

To obtain AUX1 for cryo-EM analysis or transport assay, every liter of HEK293F cells was transiently transfected with 1.5 mg AUX1 plasmids using 4 mg 40 kDa liner polyethylenimines (PEI, YEASEN) when the cell density reached 2 × 10^6^ cells per mL. The transfected cells were cultured for 60 hours before harvest.

For protein purification of AUX1 or AUX1_IL1_BRIL, cells were collected by centrifugation (4,000 g, 10 minutes) and resuspended in the lysis buffer containing 25 mM HEPES pH 7.4, 150 mM NaCl, and the Amresco protease inhibitor cocktail (2.7 μg/mL Pepstatin, 5 μg/mL Leupeptin, and 1.3 μg/mL Aprotinin). Resuspended cells were solubilized using 1% (w/v) LMNG (Anatrace) and 0.1% (w/v) CHS (Anatrace) at 4 °C overnight. After centrifugation at 12,500 r.p.m. for 1.5 hours, the supernatant was applied to anti-Flag M2 affinity resin (Sigma). The resin was rinsed using the lysis buffer supplemented with 0.005% LMNG (w/v) and 0.0005% CHS (w/v), which will be referred as buffer-W. Target proteins were eluted using buffer-W plus 200 ug/mL FLAG peptide. The eluent was then passed through the Ni-NTA resin (Qiagen) and rinsed using buffer-W supplemented with 30 mM imidazole. The target protein was eluted using buffer-W plus 300 mM imidazole. The eluent was then concentrated to approximately 0.8 mL and applied to a Superdex® 200 Increase 10/300GL column (Cytiva) that was pre-equilibrated with the running buffer containing 25 mM Tris pH 7.4 (at room temperature), 150 mM NaCl, 0.002% LMNG, and 0.0002% CHS. The peak fractions were pooled and concentrated for cryo-sample preparation. The running buffer for proteins used for functional assay contains 25 mM HEPES pH 7.4, 150 mM NaCl, 0.002% LMNG, and 0.0002% CHS.

For purification of anti-BRIL Fab and anti-Fab nanobody, the cell medium was harvested and concentrated using a 10-kDa filter (P2C010C01, Pellicon). The concentrated solution was mixed with Ni-NTA resin, washed with 10 mL wash buffer (25 mM Tris pH 7.4, 150 mM NaCl, and 30 mM imidazole) for three times, and eluted using 10 mL elution buffer (25 mM Tris pH 7.4, 150 mM NaCl, and 300 mM imidazole). The eluted protein was further purified through a Superdex® 200 Increase 10/300GL column (Cytiva) with running buffer containing 25 mM Tris pH 7.4, and 150 mM NaCl).

### [^3^H]IAA transport assay

The WT or mutant AUX1 with an N-terminal EGFP tag was subcloned into the pCAG vector. HEK293T cells at a density of about 1.0 × 10^6^ were cultured in 48-well plates. When the cell confluence reached ∼ 60%, 1.5 μg plasmids for WT or mutant AUX1 were transiently transfected into HEK293T cells using PEI. Empty vector was transfected as the control. After being cultured for 24 hours at 37 °C, cells were washed using PBS and incubated with 120 μL MEM medium pH 6.0 (iCell) containing about 30 nM radiolabeled [^3^H]IAA (specific activity 20 Ci mmol^−1^, American Radiolabeled Chemicals) at 37 °C for 10 minutes. For the transport inhibition assay, each inhibitor at a final concentration of 50 μM was added into the MEM medium. Cells were then washed three times with ice-cold Ringer’s buffer (10 mM HEPES-NaOH, 1 mM NaHCO_3_, 115 mM NaCl, 2.5 mM KCl, 1 mM MgCl_2_, 1.8 mM CaCl_2_) pH 6.4 supplemented with 10 mM unlabeled IAA (Sigma). Cells were then lysed using 150 μL 2% (w/v) SDS and mixed with 600 μL liquid scintillation. The radioactivity was measured using MicroBeta2 Microplate counter (PerkinElmer). The readout was processed using Prism 9.5.1.

To monitor the protein expression level of WT or mutant AUX1 in the [^3^H]IAA assay, EGFP fluoresce was examined using flow cytometry with the following protocol. HEK293T cells were cultured to ∼ 60% confluency in the 48-well plates and transiently transfected with 1.5 μg target plasmids per well using PEI. After culturing for 48 hours at 37 °C, cells were washed twice using PBS and resuspended in PBS buffer supplemented with 2% (v/v) fetal bovine serum (FBS). Cells were then filtered with 200 mesh nylon filter. EGFP fluorescence was detected at 488 nm by Bio-Rad ZE5 (Bio-Rad). Data analysis was performed using FlowJo 10.4.

### Cryo-EM sample preparation and data acquisition

To obtain the cryo-EM structure of the AUX1-IAA complex, purified AUX1 was concentrated to approximately 9 mg/mL and incubated with excess IAA on ice for 1 hour. For cryo-EM analysis of IAA-free AUX1, purified AUX1_IL1_BRIL, anti-BRIL Fab, and anti-Fab nanobody were incubated on ice for 1 hour with a molar ratio of approximately 1:1.4:2. Then the mixture was applied to a Superose® 6 Increase 10/300GL column (Cytiva) equilibrated with 25 mM HEPES pH 7.4, 150 mM NaCl, 0.002% LMNG, and 0.0002% CHS. Peak fractions corresponding to the ternary complex were collected and concentrated to approximately 7 mg/mL. For cryo-sample preparation, holey carbon grids (Quantifoil, Au, 300 mesh, R1.2/1.3) were subjected to glow-discharge in a vacuum for 26 seconds with mild force using the Plasma Cleaner PDC-32G-2 (Harrick Plasma Company). Aliquots of 3.5 μL were applied to the grids, blotted for 4 seconds under 100% humidity at 8 °C, and then plunged into liquid ethane cooled by liquid nitrogen using Vitrobot Mark IV (Thermo Fisher Scientific).

The prepared grids were loaded into a Titan Krios electron microscope (Thermo Fisher Scientific) equipped with a GIF Quantum energy filter (20 eV slit width) and a Gatan K3 Summit direct electron detector, operated at 300 kV. Images were auto-collected using EPU (Thermo Fisher Scientific) following standard operation procedures. A nominal magnification of ×81,000 was used for image acquisition, resulting in a pixel size of 0.53865 Å. The defocus values were set from -1.4 to -2.0 μm or -1.4 to -1.8 μm. Each micrograph was acquired with an exposure time of 5.6 s and a total electron dose of 50 e^−^/Å^2^ dose fractionated to 32 frames. Motion correction was performed using MotionCor2 (2) with a binning factor of 2, generating a pixel size of 1.0773 Å. Dose weighting was simultaneously applied, and the defocus value was calculated with Gctf (Zhang, 2016).

### Image processing

All data processing was performed using cryoSPARC (3). A flowchart that illustrates the EM data processing of the AUX1-IAA complex can be found in Figure S2D. To be brief, 3,837 high-quality micrographs were selected for preliminary data processing. A total of 6,807,488 picked particles were extracted and subjected to 2D classification. After several rounds of 2D classification, 615,674 good particles were selected for Ab_initio 3D reconstruction, yielding a good initial map from 203,593 particles. This map was then used as a template for further data processing. An additional 10,229 micrographs were collected, yielding a total of 54,584,571 particles. After 2D classification, 4,002,124 particles were selected for multiple rounds of Ab_initio 3D reconstruction. The particles from all good classes were merged and duplicates were removed, leaving 2,001,500 particles for three rounds of heterogeneous refinement. Then 351,874 particles were selected for 2D classification or 3D classification, resulting in refined maps at 4.19 Å (173,640 particles) and 3.9 Å (131,289 particles) through non-uniform refinement. The 3.9 Å map was further processed by local refinement, achieving a final map with an overall resolution of 3.5 Å. Local resolution was estimated using UCSF ChimeraX (4, 5).

A similar approach was applied for data processing of the AUX1_IL1_BRIL/Fab/Nb complex. A flowchart for the EM data processing is shown in Figure S8C. In sum, the data processing started with 20,418 micrographs. After two rounds of template picking, 2D classification, heterogeneous refinement, and Ab_initio 3D reconstruction, a 3.97 Å map was obtained (Figure S8C-8F).

### Model building and refinement

De novo model building was performed in Coot (6-8) using the 3.5 Å map of AUX1. The atomic model of AUX1 was initially constructed based on the structure predicted using AlphaFold2 (9, 10). The predicted model was manually docked into the AUX1 map using UCSF ChimeraX. Residue assignment was achieved based on the well-defined map in Coot. The model was further improved through iterative manual adjustments in Coot, followed by real_space_refinement in Phenix (11, 12). The coordination of IAA was modeled using the ligand builder eLBOW (13). To avoid overfitting, the entire model was refined against one of the two independent maps from the gold-standard refinement approach, and validation was done using the refined model (11, 12). The final model comprises 415 residues. The model of AUX1_IL1_BRIL in complex with anti-BRIL Fab and Fab-specific nanobody (Nb) was built using the model of AUX1 as a template. Statistics for the 3D map reconstruction and model refinement were provided in Table S1. All figures were prepared using UCSF ChimeraX and Coot (6-8).

### Surface plasmon resonance (SPR) assay

A Biacore 1K (Cytiva) system was employed to measure the binding affinity between AUX1 and IAA. The purified protein, diluted to 5 ug/ml with sodium acetate buffer (pH 5.0), was immobilized onto the CM5 sensor chip (Cytiva) by amine coupling to achieve approximate target densities of ∼10,000 resonance unit (RU). IAA at indicated concentrations was flowed over the chip surface at a rate of 30 μL/min with a contact time of 400 seconds. A blank control was performed using one channel of the sensor chip to exclude unspecific binding responses. The running buffer contained 150 mM NaCl, 0.01% LMNG, and 0.001% CHS, with 25 mM HEPES pH 7.4, or 25 mM MES pH 5.5. Data analysis was done using the Biacore™ Insight Evaluation Software. The results were fitted using a steady-state affinity binding model.

### All-atom molecular dynamics simulation

AUX1 was embedded into a POPC lipid bilayer using the CHARMM-GUI software package (14). Each system was solvated using the TIP3P water model, and 150 mM Na^+^ and Cl^-^ were added to neutralize the system. A periodic rectangular box (∼10 nm × 10 nm × 11 nm) was applied, containing approximately 105,000 atoms. The CHARMM36m force field was employed for proteins and lipids (15), and IAA parameters were generated via the CHARMM-GUI Ligand Reader module (16). System preparation began with 50,000 steps of energy minimization using the steepest descent algorithm, followed by a multi-phase equilibration under both isothermal–isovolumetric (NVT) and isothermal–isobaric (NPT) conditions, during which positional restraints were gradually relaxed. Unrestrained production simulations were performed using OpenMM (17), with each simulation run for 1 μs using a 2 fs integration time step at 300 K. To ensure statistical reliability, three independent replicas with different initial velocity seeds were generated for each of the six systems, totaling 18 simulations. Trajectory analyses were conducted using Visual Molecular Dynamics (VMD) (18) for structural visualization and quantitative evaluation of root mean square deviation (RMSD).

## Data availability

The atomic coordinates have been deposited in the Protein Data Bank with the accession code 9LVB and 9LVA for the IAA-free and IAA-bound AUX1, respectively. The cryo-EM maps have been deposited in EMDB with the accession code EMD-63418 for the IAA-free and EMD-63417 for the IAA-bound state, respectively.

## Contributions

D.J. conceived the project. D.J. designed and conducted the experiments. D. J. carried out the structure determination. F. K. carried out the model building. X. L and J. H. carried out the molecular dynamic simulations. All authors contributed to data analysis. Y. S., C.W. and D. J. wrote the manuscript.

## Acknowledgements

This work was supported by funds from the Key R&D Program of Zhejiang Province (2020C04001 to Y. S.), and Start-up funds from Westlake University. The authors thank X. Wang, Y. Zhang, Z. Jiang, Y. Gu, L. Huang, H. Chen, and R. Feng of the Westlake University Cryo-EM facility for technical assistance; Z. Liu of the Westlake University Supercomputer Center for computation support; P. Zhang and J. Liu of the Westlake University Crystallography Platform for technical support; X. Li of the Westlake University Isotopic laboratory for support; M. Liao and G. Fang of the Westlake University Microscopy facility; B. Shen and S. Ye of the Westlake University Flow Cytometry facility for technical support. The authors also thank J. Zhou for giving some support.

## Competing interests

The authors declare no competing interests.

## Notes

### Competing Interest Statement

The authors have declared no competing interest.

