## Supplemental Table 1 for "Structural basis of auxin binding and transport by *Arabidopsis thaliana* AUX1"

**This PDF file includes:**

Figures S1 to S14
Table S1
SI References

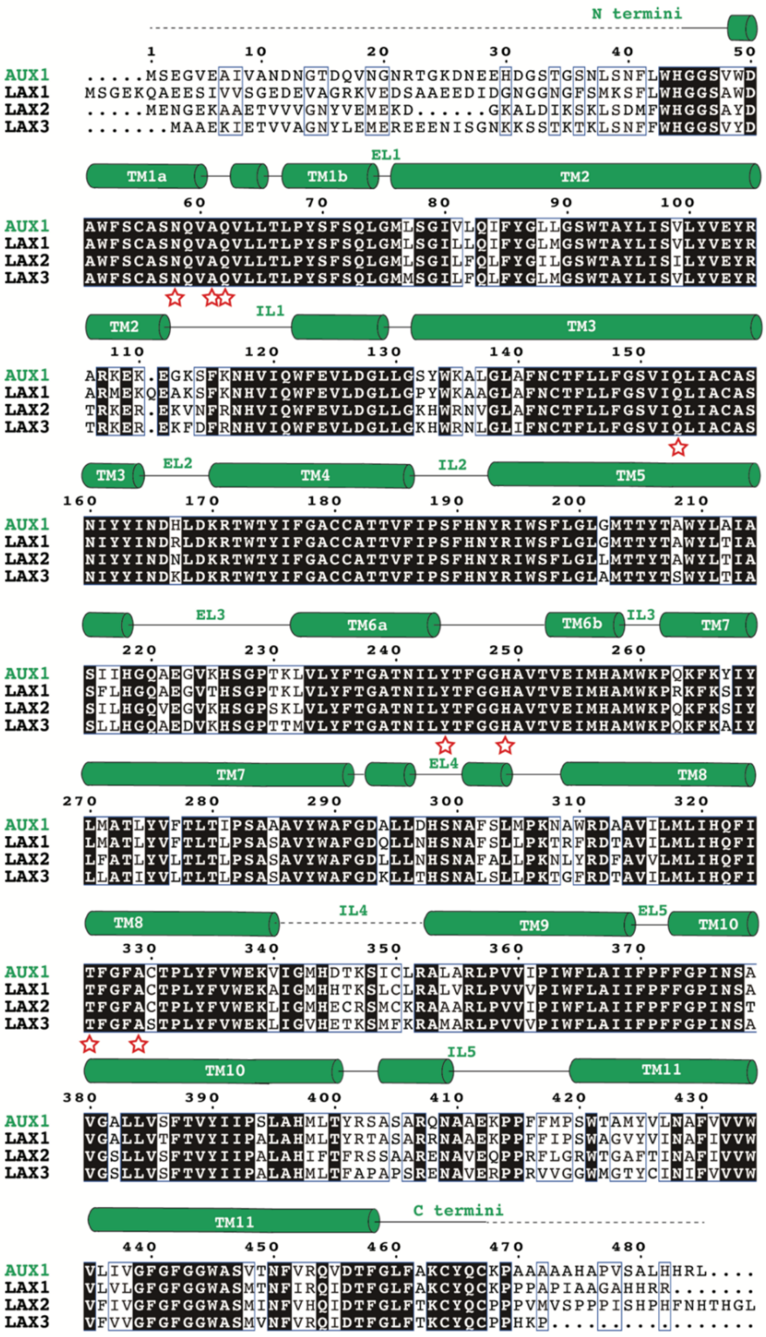


**Fig. S1. Sequence alignment of AUX1 with LAX1-3 from *Arabidopsis thaliana*.**

The sequences were aligned using CLUSTALW (1) (http://www.genome.jp/tools-bin/clustalw). Invariant residues are shaded black, and conserved residues are boxed by blue lines. Secondary structural elements of AUX1 are depicted above the sequences. The residues involved in IAA binding are indicated by red pentagons.

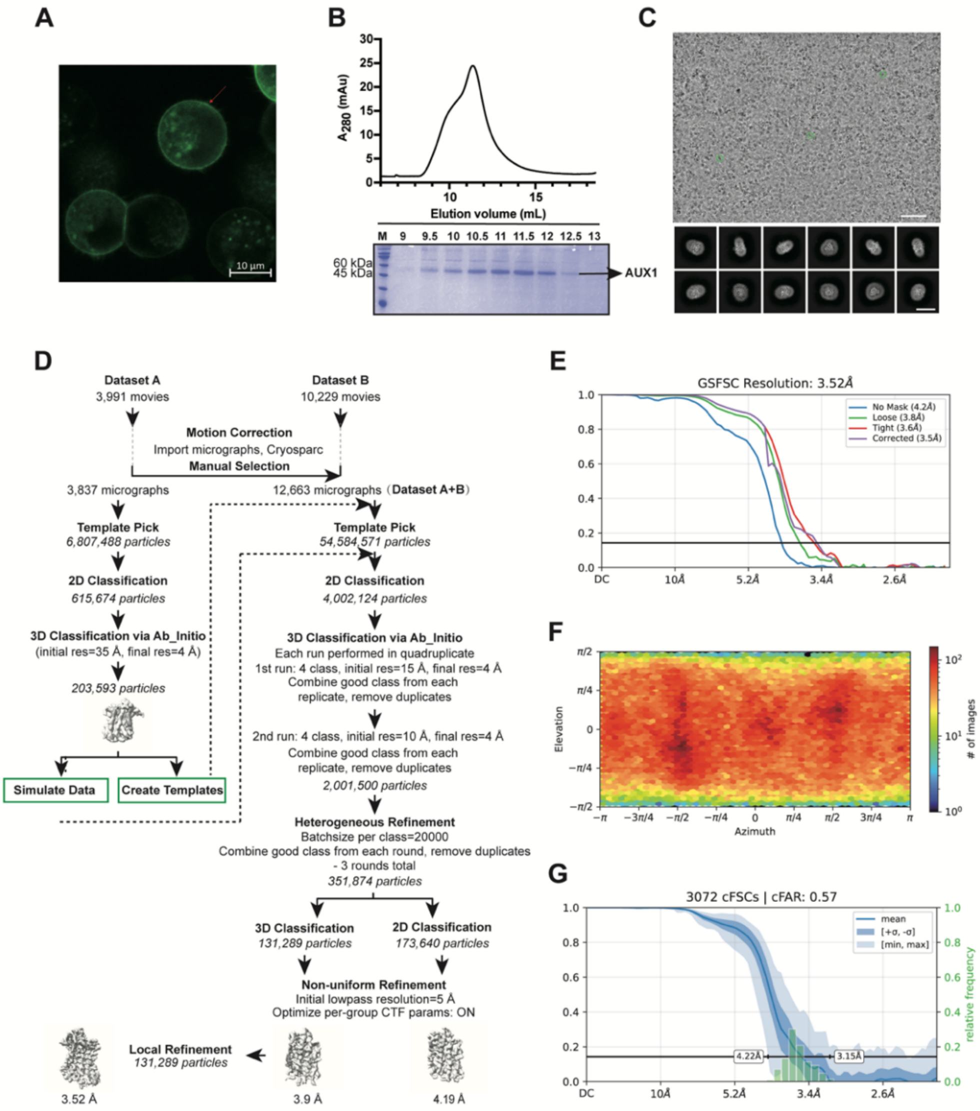

**Fig. S2. Expression, purification, and cryo-EM structural determination of IAA-bound AUX1.**

**(A)** A confocal image of HEK293F cells expressing the EGFP_AUX1 fusion protein (green).

**(B)** Purification of the AUX1 by size-exclusion chromatogram (SEC). The peak fractions were applied to SDS-PAGE and visualized by Coomassie blue staining.

**(C)** A representative cryo-EM image and 2D class averages. The scale bar in the micrograph is 90 nm.

**(D)** A flowchart of cryo-EM data processing. Details can be found in the Methods.

**(E)** The gold standard Fourier Shell Correlation (GSFSC) curves for the final 3D refinement.

**(F)** The angular distribution plot of the particles in the final 3D reconstruction.

**(G)** Corrected Fourier Shell Correlation (cFSC) plot of spatial frequency.

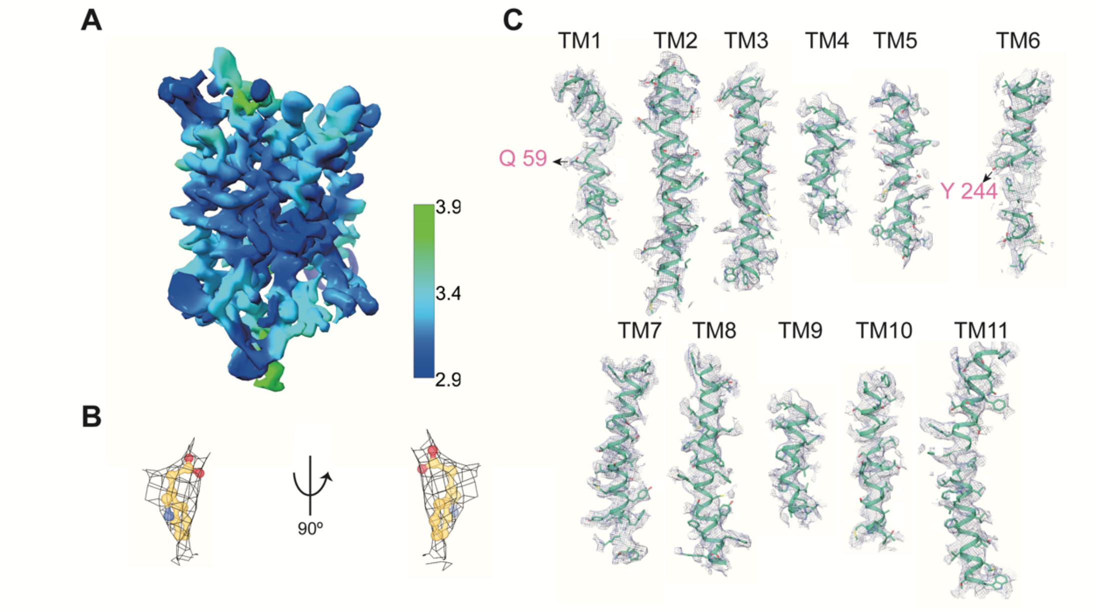


**Fig. S3. Representative EM densities of IAA-bound AUX1.**

**(A)** The EM density map is colored by local resolutions. Local resolution is calculated using cryoCPARC and represented in UCSF ChimeraX.

**(B)** The EM density of IAA was generated in UCSF ChimeraX at the contour level of 0.86.

**(C)** EM maps of the 11 transmembrane segments of AUX1. The densities, shown as grey meshes, of representative segments were generated in UCSF ChimeraX at the contour level of 0.5. Two residues that are involved in conformational changes of TM1a are labeled.

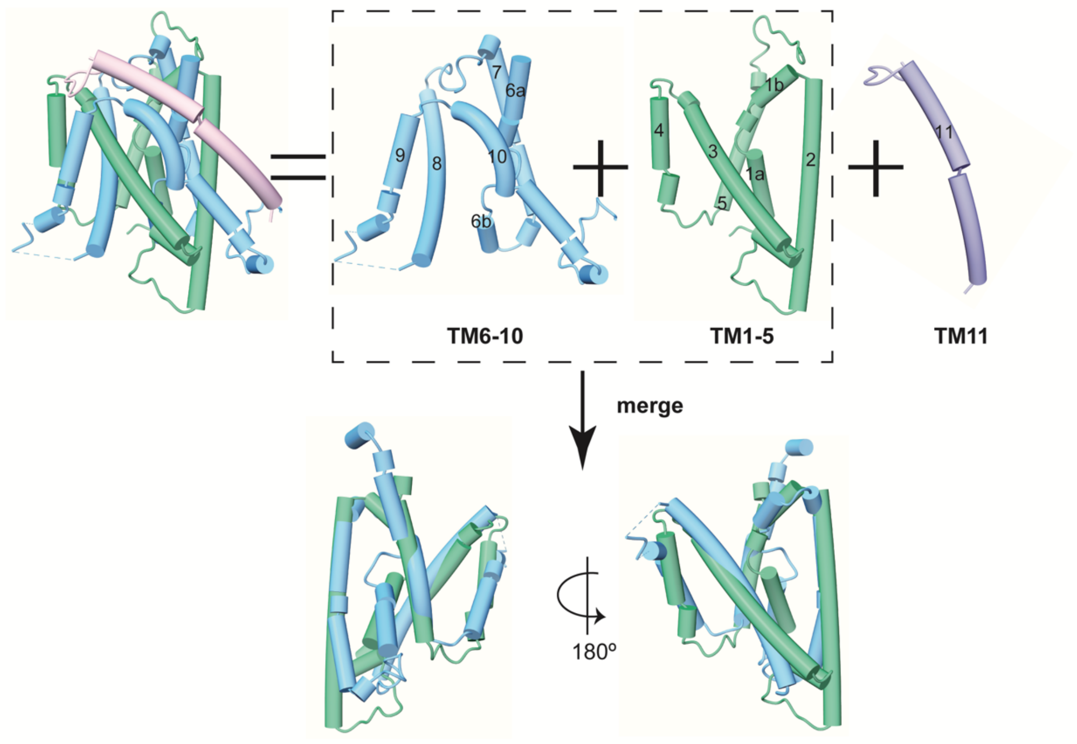


**Fig. S4. AUX1 conforms to the classic LeuT-fold.**

AUX1 has 11 transmembrane segments (TMs 1-11). TMs 1-5 (green) and TMs 6-10 (blue) of AUX1 are structurally inverted repeats.

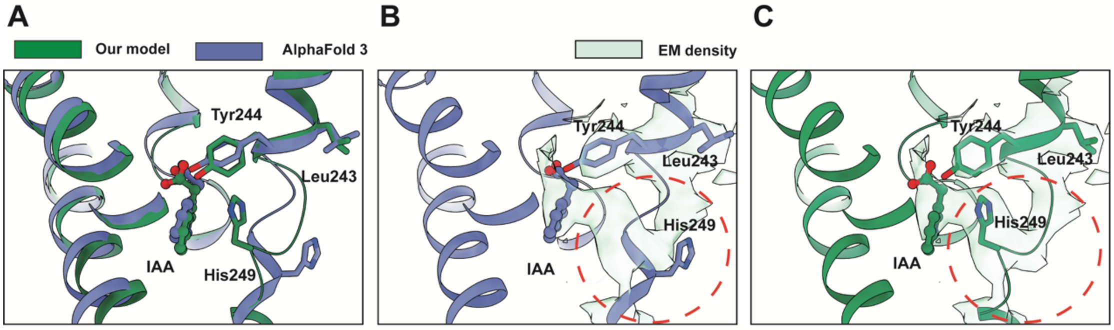


**Fig. S5. Model building of IAA binding pocket.**

**(A)** Superimposition of our IAA-bound AUX1 structure with the AlphaFold3-predicted model (2).

**(B)** The main chain around His249 in the AlphaFold3-predicted model cannot fit into our EM density. The densities are contoured at the level 1.5 in UCSF ChimeraX.

**(C)** Our model fits the local EM map in the IAA binding pocket.

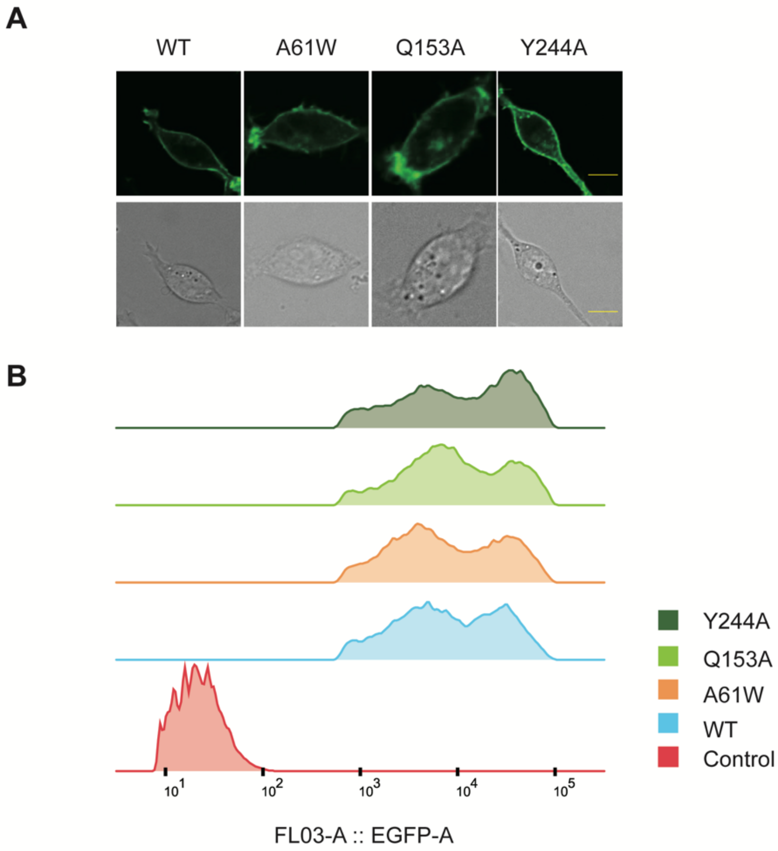


**Fig. S6. Protein expression of WT and mutant AUX1.**

**(A)** Representative fluorescent (upper) and bright-field (lower) images of cells expressing green fluorescent protein (EGFP)-tagged WT (3) and mutant AUX1.

**(B)** Flow cytometry assessment of protein expression levels of the WT and mutant AUX1 proteins.

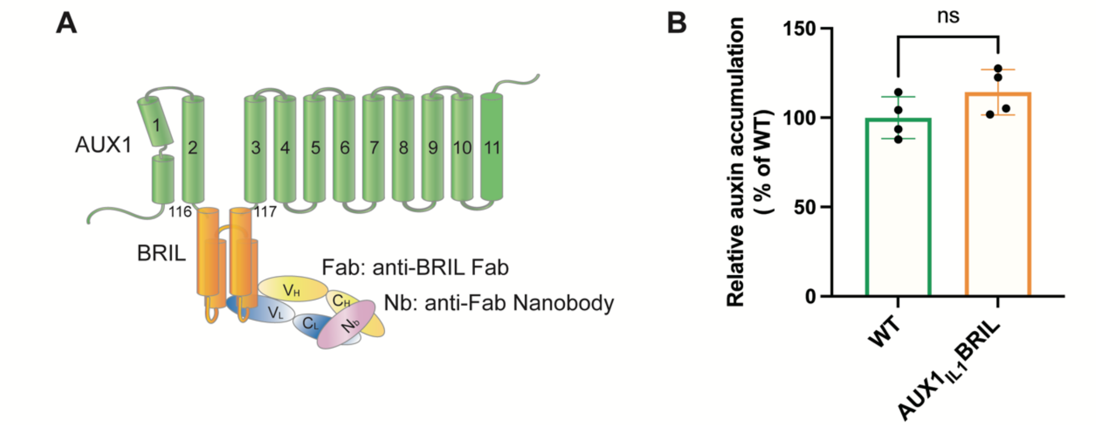


**Fig. S7. Engineering of AUX1 for structural determination of the IAA-free transporter.**

**(A)** Schematic diagram of the $AUX1_{IL1}$BRIL/Fab/Nb complex. A BRIL was inserted in the loop between TM2 and TM3, defined as IL1 or the intracellular loop 1.

**(B)** $AUX1_{IL1}$BRIL maintains the same transport activity as the WT AUX1. [$^3$H]IAA uptake assay was performed to verify the activity of the fusion protein. The results are normalized to the activity of WT AUX1. Data are mean ± s.d.; n=4 (ns, non-significant, Two-tailed unpaired t-test). All experiments were repeated for three times.

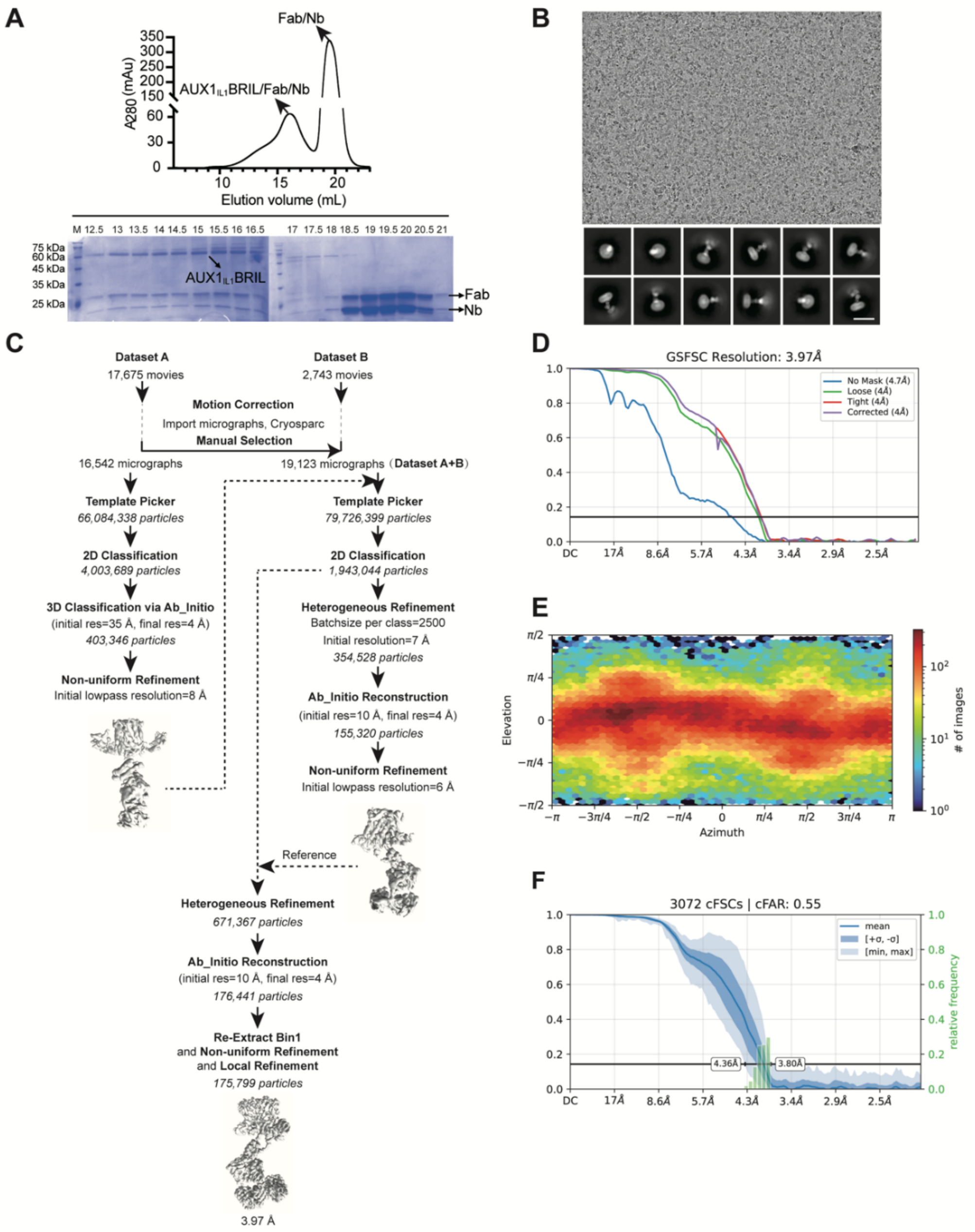


**Fig. S8. Structure determination of AUX1$_{IL1}$BRIL/Fab/Nb complex.**

**(A)** Purification of AUX1$_{IL1}$BRIL complexed with Fab and anti-Fab nanobody by size-exclusion chromatogram (SEC). The peak fractions from gel filtration were applied to SDS-PAGE and visualized by Coomassie blue staining.

**(B)** A representative EM micrograph and 2D class averages. The scale bar is 90 nm.

**(C)** A flowchart of cryo-EM data processing for two datasets using cryoSPARC.

**(D)** The gold standard Fourier Shell Correlation (GSFSC) curves for the final 3D refinement.

**(E)** The angular distribution plot of the particles in the final 3D reconstruction.

**(F)** Corrected Fourier Shell Correlation (cFSC) plot of spatial frequency.

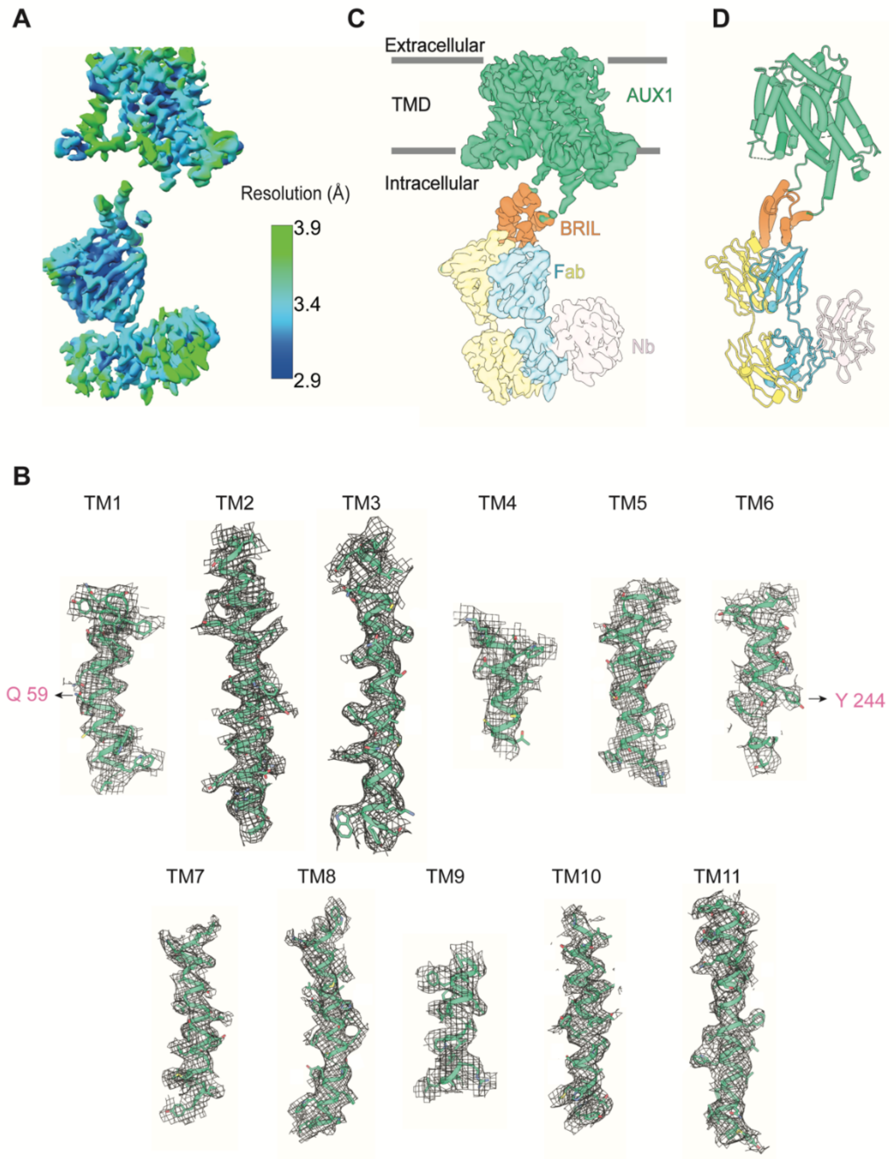


**Fig. S9. Cryo-EM analysis of the AUX1$_{IL1}$BRIL/Fab/Nb complex.**

**(A)** EM reconstruction of the AUX1$_{IL1}$BRIL/Fab/Nb complex. Color-coded local resolutions were calculated in cryoSPARC; the map was generated in UCSF ChimeraX (4, 5).

**(B)** EM maps of the 11 transmembrane segments. The densities are contoured at the level 0.5 in UCSF ChimeraX. Two residues that are involved in conformational changes of TM1a are labeled.

**(C)** Cryo-EM map of the AUX1$_{IL1}$BRIL/Fab/Nb complex. The map was contoured at the level 0.5 in UCSF ChimeraX.

**(D)** Overall structure of the AUX1$_{IL1}$BRIL/Fab/Nb complex. Each individual component is colored the same as in panel C.

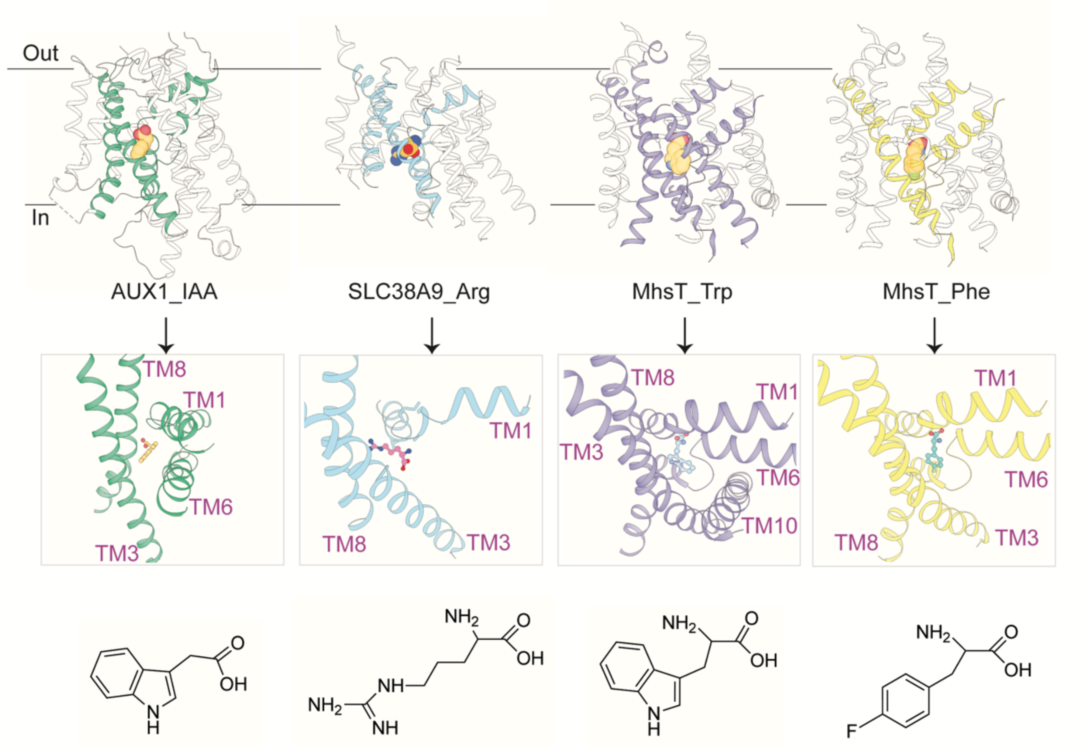


**Fig. S10. Comparison of the substrate-binding pocket between AUX1 and other AAAP subfamily transporters.**

The amino acid/auxin permease (AAAP) family members AUX1, SLC38A9, and MhsT all share the same LeuT-fold. Comparison of their structures reveals similar substrate binding sites but important variations in the amino acid residues and TMs involved. The upper panel shows the structures of AUX1 bound to IAA, SLC38A9 bound to arginine (6) (PDB code: 6C08), MhsT bound to tryptophan (PDB code: 4US3) or 4-fluoro-L-phenylalanine (PDB code: 6YU4) (7). TMs involved in substrate binding are highlighted in different colors. The lower panel shows the chemical structures of IAA, arginine, tryptophan, and 4-fluoro-L-phenylalanine. Although tryptophan serves as the biosynthetic precursor of IAA, the details for their recognition by the corresponding transporters are different.

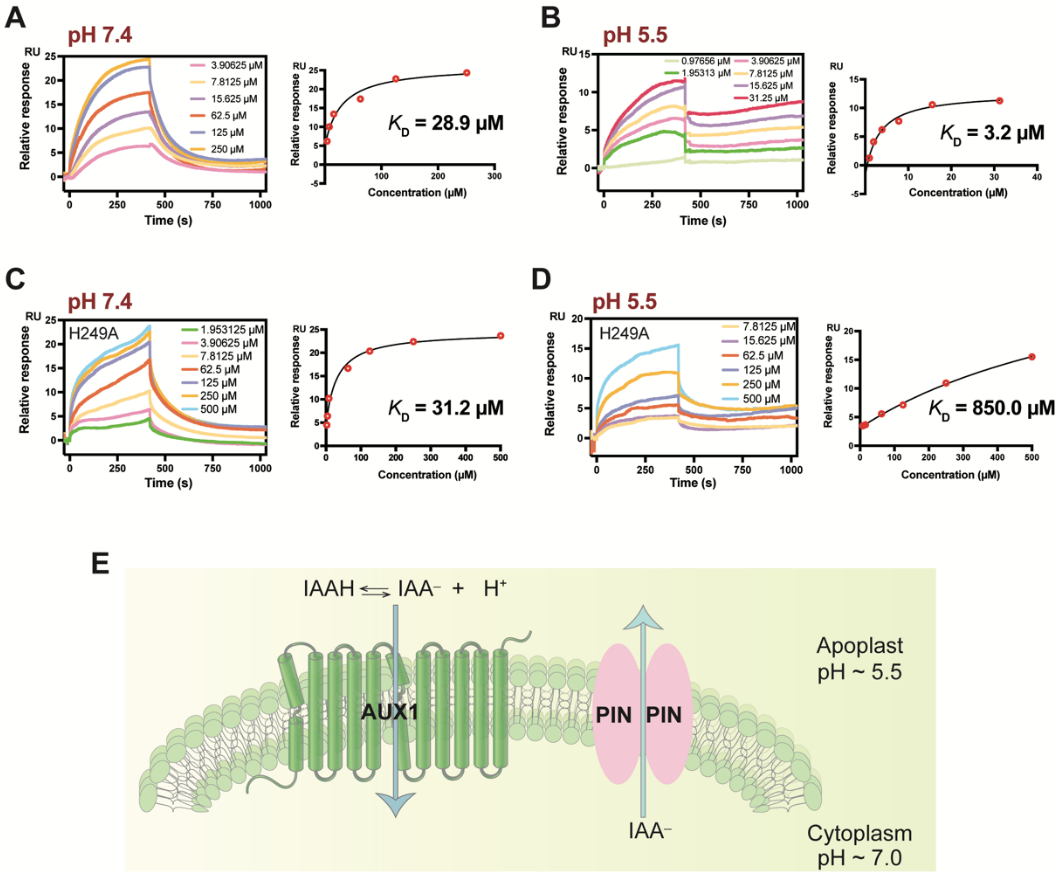


**Fig. S11. AUX1 binds to IAA with a much higher affinity at pH5.5 than at pH 7.4.**

**(A** and **B)** AUX1 binds to IAA with a higher affinity at lower pH. The binding affinities were measured using SPR. IAA has a pKa of ~ 4.75. At pH 7.4, IAA carries a negative charge at the carboxylate group. At pH 5.5, however, a small portion of IAA carboxylate becomes neutral and as a consequence may alter its interaction with AUX1. These changes may give rise to some non-reversible binding, particularly at high IAA concentrations. RU, response unit.

**(C** and **D)** The AUX1 mutant H249A binds to IAA with a much lower affinity at lower pH. The binding affinities were measured using SPR. RU, response unit.

**(E)** A cartoon model depicting the transport mechanisms of AUX1 and PIN. Transport directions through each transporter are indicated by light blue arrows.

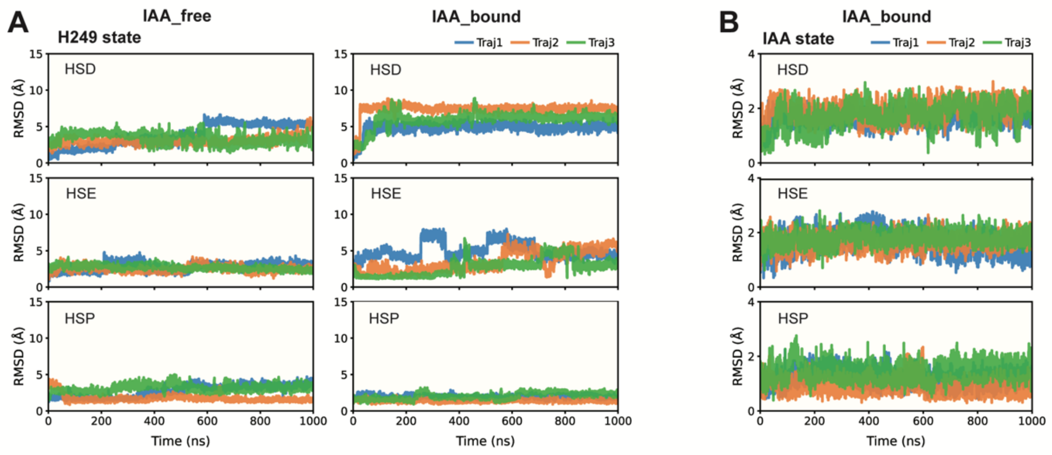


**Fig. S12. Conformational dynamics of His249 and IAA in different His249 protonation states.**

**(A)** Conformational dynamics of His249. RMSD time series of His249 in IAA_free (8) and IAA_bound AUX1 systems, comparing three protonation states (HSD, HSE, and HSP).

**(B)** Conformational dynamics of IAA in all three His249 protonation states.

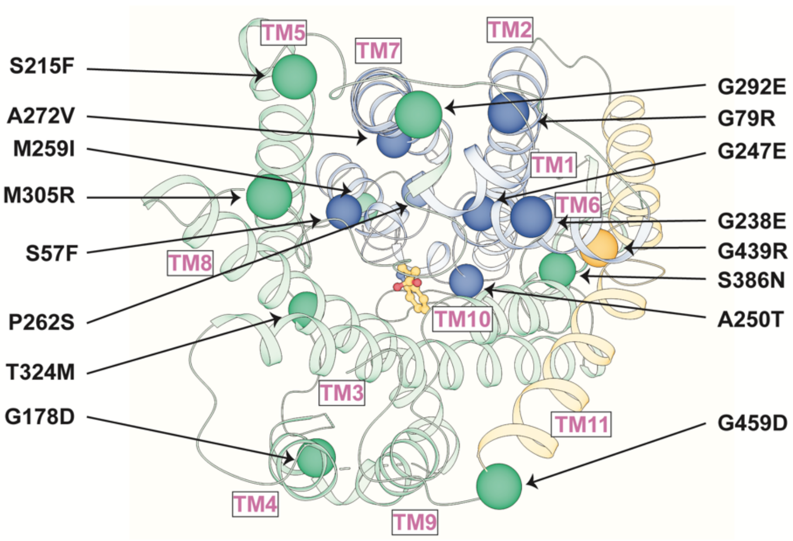


**Fig. S13. Mapping of functional mutations onto the structure of the AUX1-IAA complex.**

The core domain, scaffold domain, and TM11 are colored blue, green, and tangerine, respectively. Residues that are targeted by missense mutations are shown as spheres.

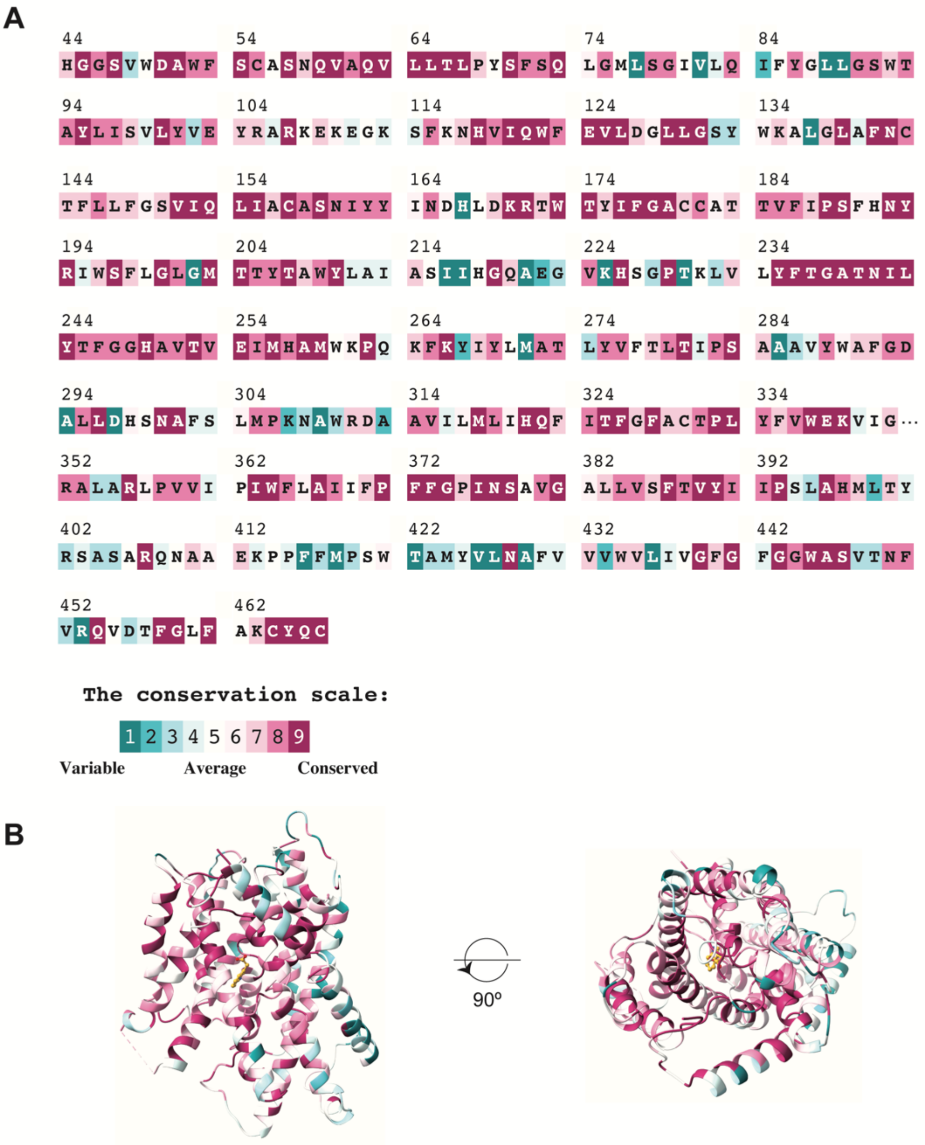


**Fig. S14. Sequence conservation of AUX1.**

**(A)** The amino acid sequences of AUX1 from *Arabidopsis thaliana*. Each amino acid is colored-coded on the basis of its degree of conservation using Consurf (9).

**(B)** The EM structure of AUX1 from *Arabidopsis thaliana* is displayed by its sequence conservation. Two perpendicular views are shown.

**Table S1. Statistics of cryo-EM data collection and model refinement**

| | *At*AUX1 | *At*AUX1-IAA |
|---|---|---|
| **Data collection** | | |
| Microscope | FEI Titan Krios | FEI Titan Krios |
| Magnification | 81,000 | 81,000 |
| Voltage | 300 | 300 |
| Electron exposure ($e^{-}$/Å$^{2}$) | 50 | 50 |
| Defocus range (μm) | -1.4 ~ -1.8 | -1.4 ~ -2.0 |
| Pixel size (Å) | 1.0773 | 1.0773 |
| Symmetry imposed | C1 | C1 |
| Raw movies | 20,418 | 12,663 |
| **Reconstruction** | | |
| Particle Number | 175,799 | 131,289 |
| Map resolution (Å) | 3.97 | 3.52 |
| FSC threshold | 0.143 | 0.143 |
| Map resolution range (Å) | | |
| **Refinement** | | |
| Protein residues | 942 | 415 |
| Ligand | --- | IAA |
| *B* factors (Å$^{2}$) | | |
| Protein | 88.19 | 75.58 |
| Ligand | --- | 62.00 |
| Water | --- | --- |
| R.m.s. deviations | | |
| Bond lengths (Å) | 0.003 | 0.002 |
| Bond angles (°) | 0.548 | 0.495 |
| **Validation** | | |
| MolProbity score | 2.24 | 2.02 |
| Clashscore | 9.73 | 8.31 |
| Ramachandran plot | | |
| Favored (%) | 94.21 | 96.59 |
| Allowed (%) | 5.79 | 3.41 |
| Disallowed (%) | 0.00 | 0.00 |
| **PDB code** | | |
| **EMDB code** | | |
